# *Lactobacillus paracasei* CNCM I-5220-derived postbiotic counteracts skin inflammation and protects skin barrier integrity

**DOI:** 10.1101/2024.11.16.623921

**Authors:** Silvia Pimazzoni, Francesca Algieri, Nina Tanaskovic, Daniele Braga, Giuseppe Penna, Maria Rescigno

**Affiliations:** Department of Biomedical Sciences, Humanitas University, 20072 Pieve Emanuele, Italy; Postbiotica S.r.l., 20123 Milan, Italy; IRCCS Humanitas Research Hospital, 20089 Rozzano, Italy

**Author notes:** These authors have contributed equally to this work (Silvia Pimazzoni, Francesca Algieri).

**Keywords:** Postbiotic, *Lactobacillus paracasei*, Skin inflammation, Skin barrier integrity, Filaggrin, Atopic dermatitis

## Abstract

**Introduction:** Skin inflammation and damage of skin barrier integrity contribute to the pathogenesis of inflammatory skin diseases such as psoriasis and atopic dermatitis. Nowadays, there is a growing interest in alternative and complementary strategies to counteract skin inflammation and treat dermatologic conditions.

**Objectives:** Postbiotics, metabolites released during bacterial fermentation, exert anti-inflammatory proprieties and contribute to the maintenance of epithelial barrier integrity. Therefore, we investigated the effect of LP-PBL, a novel *Lactobacillus paracasei* CNCM I-5220-derived postbiotic, in controlling skin inflammation and protecting the skin barrier.

**Methods:** We performed RNA-sequencing on Poly(I:C)-stimulated keratinocytes and investigated LP-PBL efficacy in regulating pro-inflammatory cytokine release. The production of most relevant cytokines was demonstrated by ELISA. Then, we tested postbiotic lenitive efficacy on healthy volunteers’ irritated skin. We compared the effect of a postbiotic-containing cream formulation versus placebo on sodium lauryl sulphate treated skin.

**Results:** We demonstrate that LP-PBL has anti-inflammatory activity by modulating Poly(I:C)-dependent inflammatory pathways and pro-inflammatory cytokine release in keratinocytes. The postbiotic inhibited the upregulation of interleukin (*IL)-23A*, which is overexpressed in psoriatic skin, and of *IL-33* and thymic stromal lymphopoietin (*TSLP*) that are upregulated in atopic dermatitis. Moreover, it increased filaggrin and zonula occludens (*ZO*)-1 expression in Poly(I:C)-stimulated keratinocytes, suggesting beneficial effects on inflamed and damaged skin. Consistently, a clinical test on healthy volunteers showed that topic LP-PBL treatment significantly reduced skin redness upon sodium lauryl sulphate challenge compared to placebo, leading to a rapid recovery of the irritated skin.

**Conclusion:** Overall, we demonstrated the protective role of LP-PBL on the skin, both *in vitro* and in the clinical test, suggesting that postbiotic topical application could represent an innovative and promising strategy to counteract skin inflammation and preserve skin barrier integrity.

## INTRODUCTION

The outer layer of the skin, the epidermis, represents the primary interface between our body and the external environment. Epidermal keratinocytes play a crucial role in maintaining the integrity and hydration of the barrier as well as protecting the skin from pathogen invasion, ultraviolet radiation, and excessive trans-epidermal water loss (TEWL) [1]. Moreover, they play a role in regulating skin immune responses: through their pattern recognition receptors (PRRs), keratinocytes can recognize different pathogen-associated molecular patterns and secrete, as well as respond, to cytokines, chemokines, and growth factors. Of the different PRRs, keratinocytes express toll-like receptor (TLR)-3 that binds viral double-stranded RNA (dsRNA) and induces the release of several cytokines and chemokines [2], some of which contribute to the onset of inflammatory skin diseases such as atopic dermatitis (AD) and psoriasis.

These pathologies are also characterized by a disrupted epidermal barrier. Indeed, AD skin shows reduced expression of epidermal lipids, natural moisturizing factors (NMFs), tight junction (TJs) proteins, and epidermal differentiation markers such as filaggrin and loricrin [3].

Furthermore, the pathogenesis of AD and psoriasis is marked by alterations in the skin microbiota, including qualitative and quantitative changes such as a significant decrease in bacterial diversity [4, 5]. Commensal bacteria, and generally the skin microbiota, play a pivotal role on the skin by directly competing with microbial pathogens for colonization and by promoting skin barrier functions, positively acting on the host immune system [6]. Therefore, the reduction of commensal bacteria abundance or changes in microbiota composition (dysbiosis) can directly contribute to the development of skin disorders [7].

First-line treatment for AD consists of anti-inflammatory drugs, which, however, exhibit limited efficacy in moderate-to-severe AD and do not counteract systemic inflammation when administered topically; while corticosteroids and immunosuppressants cannot be administered for a long period of time because of their adverse effects [8]. A similar scenario arises for the treatment of psoriasis [9]. Therefore, the identification of innovative and effective therapeutic strategies is essential for the treatment of these disorders.

Given the crucial role of skin dysbiosis in the development of inflammatory skin diseases, different approaches have been proposed to re-establish microbial equilibrium and skin homeostasis. Among these, topical administration of probiotics has been explored [10], but it has shown some limitations that need to be addressed [11]: 1. As the therapeutic efficacy of probiotics is dependent on bacteria viability, indicating that the metabolic activity of live bacteria is essential for their functionality [12], maintenance of viability of probiotics in a topical formulation remains challenging; 2. Application of living microorganisms to an irritated skin raises safety concerns for possible bacterial translocations [13]; 3. Topical probiotics administration can lead to the development of antibiotic resistance [11]; 4. Clinical studies have revealed shortcomings, particularly during acute inflammatory responses [14, 15]. Given the variable efficacy and safety concerns associated with live probiotics [16], there is a growing interest in utilizing well-characterized postbiotics as a safer alternative, particularly in acute and chronic inflammatory conditions [17–19].

Postbiotics are described as “any factor resulting from the metabolic activity of a probiotic or any released molecule capable of conferring beneficial effects to the host in a direct or indirect way” [19, 20]. Hence, in this definition, they lack bacteria (either dead or alive) or cell wall components, and exert multiple beneficial effects, including immunomodulatory, anti-inflammatory, anti-microbial, and anti-aging activities [12, 21–23]. Due to the absence of live bacteria in postbiotics, along with their elevated safety and longer shelf life [22], the administration of postbiotics may represent a novel approach in harnessing the therapeutic potential of beneficial microbes while mitigating possible risks associated with live bacteria administration [14, 24].

Since the growing need for a safe, tolerable, and effective treatment to counteract skin inflammation and the emerging role of postbiotics on the skin, we sought to investigate, *in vitro* and *in vivo*, in human tests, the activity of LP-PBL, an innovative postbiotic derived from the fermentation of lactate by *Lactobacillus paracasei* CNCM I-5220.

## MATERIALS AND METHODS

### Cell lines

Immortalized adult human epidermal keratinocytes HaCaT were purchased from Cell Lines Service GmbH (Eppelheim, Germany) and maintained in Dulbecco’s modified Eagle’s medium (DMEM) supplemented with 10% Fetal Bovine Serum, 1% Glutamine, 1% Penicillin-Streptomycin and cultured at 37 °C with 5% CO_2_.

### Postbiotic production

Postbiotica S.r.l. generated the postbiotic LP-PBL starting from an inoculum of *L. paracasei* CNCM I-5220. Once the bacteria were grown at 37°C in the fermentation medium, they were collected to undergo the fermentation process following the PBTech® technology (described in patent WO 2019 /149940). The fermentation was carried out for 24 hours in a specific fermentation solution containing sodium lactate. Bacteria were then centrifuged (4000 rpm at 4°C) and the collected supernatant was pasteurized at 90°C for 10 minutes to ensure the removal of live bacteria. The supernatant was supplemented with mannitol and then subjected to freeze-drying.

### RNA isolation and gene expression analysis

HaCaT cells were seeded in 6-well plates at 2 x 10^5^ cells/well and, after 24 hours, were starved overnight in DMEM with 2% FBS. Cells were treated with different concentrations (0.1 and 1 mg/ml) of LP-PBL or its vehicle containing mannitol following the stimulation with 1 µg/ml Poly(I:C) (#5-pic-tlrl, InvivoGen). After 6 hours, cells were collected and lysed prior to mRNA extraction (DirectZol RNA MiniPrep Plus #R2072, Zymo Research). The extracted RNA was used as template for subsequent cDNA synthesis using ImProm-II™ Reverse Transcriptase (#A3803, Promega) and random hexamers. Real-time PCR reactions were performed with Luna Universal qPCR Master Mix (#M3003E, New England Biolabs) in QuantStudio 7 Flex Real-Time PCR System. Primers used in this study are listed in **Table 1**. The relative gene expression was quantified using the 2–ΔΔCt method and normalized to glyceraldehyde 3-phosphate dehydrogenase (*GAPDH*) mRNA. Data are presented as a fold change compared to untreated cells.

**Table 1:**
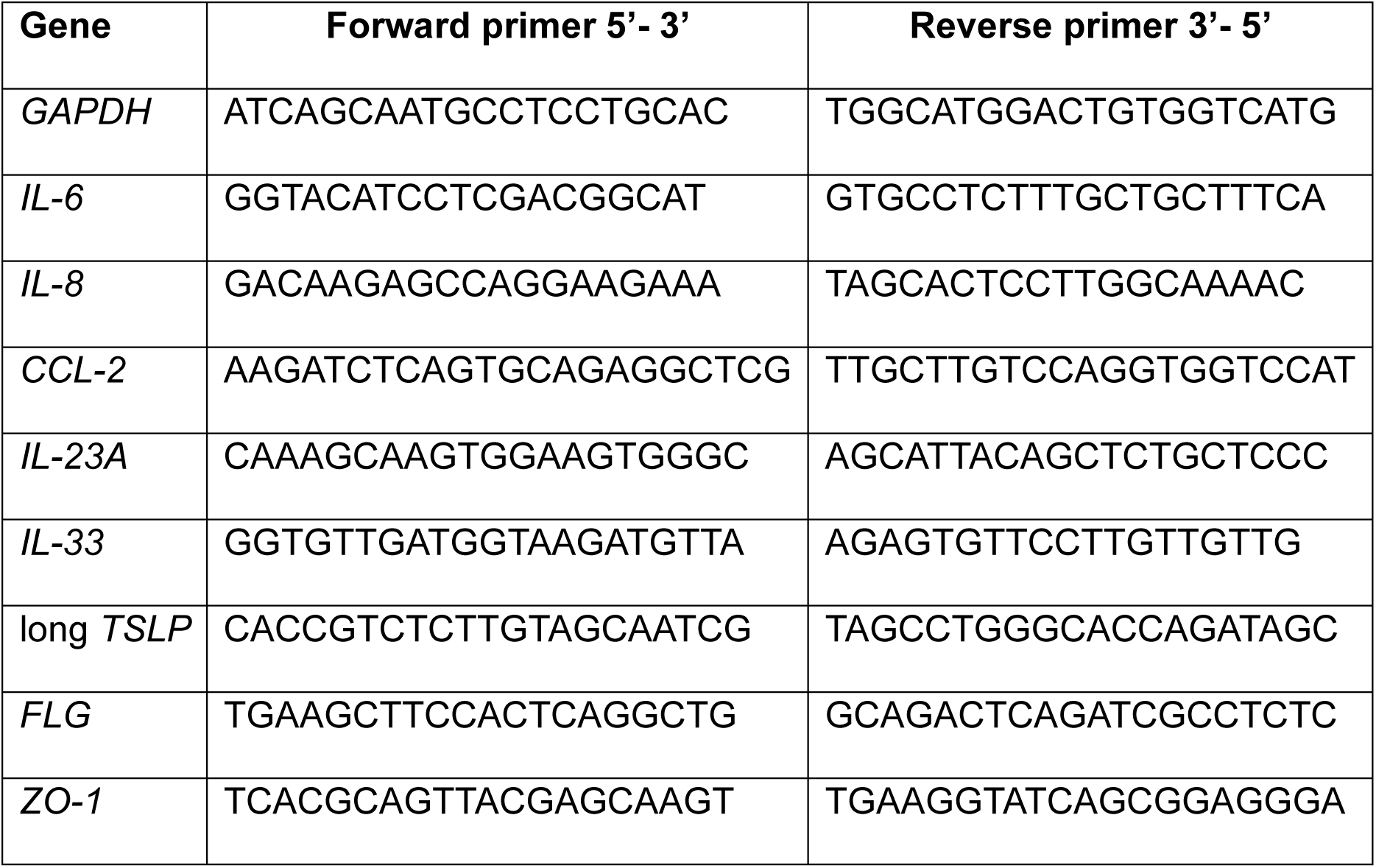
Human primers designed for real-time PCR.

### RNA QC, library preparation and sequencing

RNA quality control was performed with Agilent 4200 Tape Station system using RNA ScreenTape analysis kit (Agilent, Santa Clara, CA, USA). Only RNAs having RNA Integrity Number (RIN)>8 were used for library preparation. Libraries for mRNA sequencing were prepared with the SMART-Seq v4 PLUS Kit (Clontech-Takara). D5000 ScreenTape assay was used to check final libraries, while D5000 High Sensitivity (HS) ScreenTape assay to check pools. All samples were sequenced on an Illumina NextSeq 2000 P3 Reagents (100 Cycles, #20040559) producing on average 22.54 ± 4.48 M 76-bp single-end reads per sample.

### RNA-sequencing data analysis

Quality control of raw data was performed with fastqc v0.11.8. Quality filtering was performed with trimmomatic v0.39 setting the following parameters: CROP:74 MINLEN:50 AVGQUAL:25. High-quality reads were aligned to the human reference genome (GRCh38 primary assembly genome) using STAR (v2.7.6a). Reads were assigned to genes with featureCounts (v2.0.1) using the gencode.v42.annotation.gtf Gene Transfer File as a reference annotation file for genomic feature boundaries. DESeq2 (v1.38.3) R package was used for processing of raw counts, exploratory data analysis and differential expression analysis. Gene Set Enrichment Analysis (GSEA) was performed to investigate the pathways most modulated by the treatment. Only differentially expressed genes with following criteria were used: p_adj_< 0.05 and |log_2_ Fold Change| > 0.5. The files used as input for pre-ranked GSEA analysis were the list of differentially expressed genes of the experiment ranked by log_2_ Fold Change and the reference files h.all.v2023.2.Hs.symbols.gmt and c5.all.v2023.2.Hs.symbols.gmt. The heatmap was generated with pheatmap R package: genes of the TLR3 cascade as described in the Reactome database were selected for visualization. Normalized data obtained with DESeq2 were log_2_ transformed, given as input to the pheatmap() function and row-wise scaled.

### Enzyme-linked immunosorbent assay (ELISA)

HaCaT cells were seeded in 96-well flat bottom plates at a density of 4 x 10^4^ cells/well. 24 hours after the seeding, complete medium was replaced by starvation medium overnight. Cells were treated with different concentrations of LP-PBL, vehicle, and/or stimulated with 1 µg/ml Poly(I:C). After 24 hours, supernatants were collected for protein evaluation performed with enzyme-linked immunosorbent assay (ELISA) kit (R&D System, Inc. Bio-Techne) according to manufacturer’s instruction in order to assess human IL-6 (#DY206), IL-8 (#DY208), and CCL2 (#DY279).

### Immunofluorescence analysis

HaCaT keratinocytes were seeded in Permanox chambers (#177410, Thermo Fisher Scientific) at a density of 2 x 10^4^ cells/well and allowed to reach 70% confluence, upon which cells were maintained in overnight starvation. Then, cells were treated with 1 mg/mL LP-PBL or vehicle and/or stimulated with 1 µg/ml Poly(I:C). After 24 hours, samples were fixed in 4% paraformaldehyde, permeabilized with 0.25% Triton X-100, and then incubated with blocking solution (1% BSA in PBS). Staining was performed using primary antibody overnight at 4°C. Samples were then washed with 0.1 M Tris-HCl pH 7.4 and stained with secondary antibody for 2 hours. Nuclei were counterstained with DAPI. Sections were mounted with Vectashield mounting medium and images were acquired with Leica TCS SP8I Laser scanning confocal microscope 63X/1.30 oil immersion objective. All images were analyzed with Fiji (ImageJ) software version 2.3.0. Antibodies used for these staining: anti-ZO-1 monoclonal antibody AF488 (#339188, Invitrogen 1:100); anti-filaggrin (#ss-66192, Santa Cruz Biotechnology, 1:100); Phalloidin AF488 (#A12379, Invitrogen, 1:400); secondary antibody donkey anti-mouse IgG AF555 (#A-31570, Invitrogen, 1:400).

### MTT cytotoxicity assay

To evaluate HaCaT viability, MTT (Thiazolyl Blue Tetrazolium Bromide, #ab146345, Abcam) colorimetric assay was used. After 24 hours of postbiotic treatment, supernatants were removed and cells were incubated with 0.45 mg/ml MTT in starvation medium at 37 °C for 3 hours. Pure dimethyl sulfoxide (DMSO) was used to solubilize the formazan crystals prior to measuring the absorbance at 570 nm (reference wavelength of 640 nm) on CLARIOstar microplate reader (BMG Labtech).

### Instrumental evaluations of LP-PBL efficacy

To evaluate the lenitive efficacy of a cosmetic cream containing 2% of LP-PBL or placebo, a clinical test was carried out by Institute of Skin and Product Evaluation (ISPE s.r.l. in Milan, Italy) according to the principles of Good Laboratory Practice, Good Clinical Practice, and the Declaration of Helsinki. The clinical test was conducted on 15 healthy volunteers (13 females and 2 males) with an average age 53.9 following the inclusion and exclusion criteria listed in **Table 2**. Occlusive patches containing 2% Sodium Lauryl Sulfate (SLS) aqueous solution were applied for 24 hours on four selected areas of volunteers’ forearms using Finn Chambers. Two of the four areas were treated three times a day for two days with the cream containing LP-PBL, one with the placebo and one was left untreated. Instrumental measurements of TEWL and skin redness were taken before patches application and 24, 48, and 72 hours after patches removal. The quantification of skin redness was performed with a Chromameter CR 400 (Konica Minolta) considering only red-green color axis, while TEWL was measured by a Tewameter TM 300 MDD 4 (Courage & Khazaka).

**Table 2:**
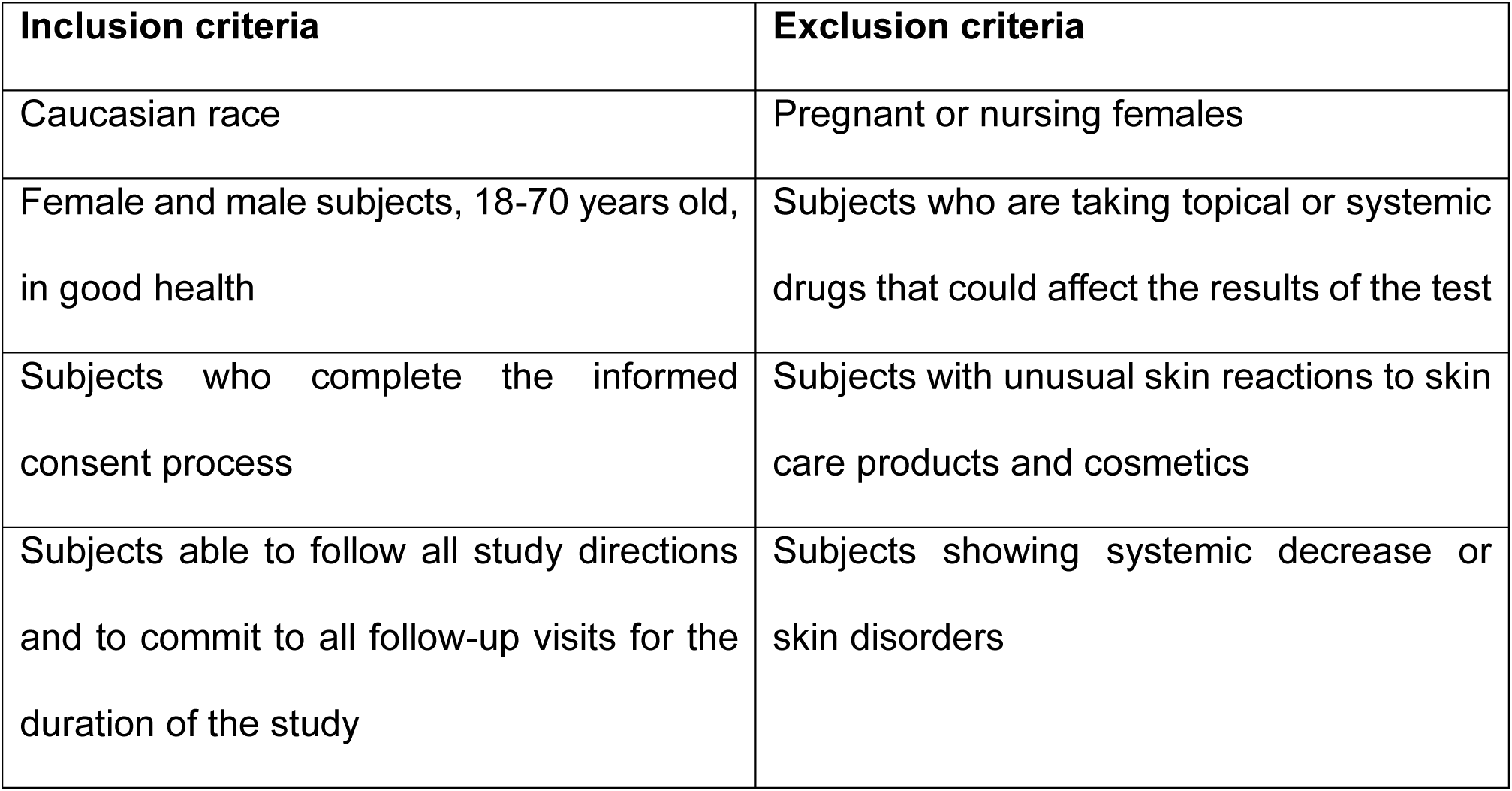
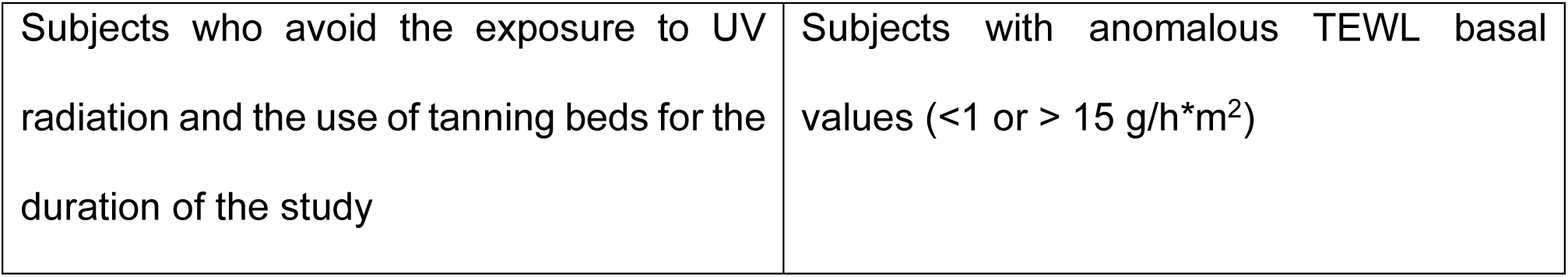
Inclusion and exclusion criteria for the clinical test.

### Statistical analyses

All the data are the results of two or three independent experiments. Statistical analyses were performed using GraphPad Prism 9.4.1 software. Data were analyzed using paired or unpaired t-test, one-way ANOVA (Tukey’s multiple comparisons test) or two-way ANOVA (Šidák multiple comparisons test). Results were represented as mean ± SEM. In the figures: *p < 0.05, **p < 0.01, ***p < 0.001, ****p<0.0001.

## RESULTS

### LP-PBL treatment shows anti-inflammatory signature on Poly(I:C)-stimulated keratinocytes

Epidermal keratinocytes are pivotal in shaping the skin barrier and modulating both innate and adaptive immune responses against external stimuli. Alterations in these responses can contribute to the onset of various skin diseases. Indeed, keratinocytes express a variety of TLRs, including TLR3 that recognizes viral dsRNA [2].

To assess whether postbiotic treatment could control cytokine and chemokine release upon TLR3 engagement, we stimulated keratinocytes with Poly(I:C), one of the most potent activators of TLR3 [25], treated them with LP-PBL or vehicle for 6 hours and performed RNA-Seq analysis. Volcano plots showed that, in steady-state, LP-PBL treatment induced only minimal changes in the transcriptomic profile compared to vehicle, while Poly(I:C)-stimulation modulated several genes (**Fig. S1**). In the presence of Poly(I:C), vehicle treatment did not influence gene expression, whereas LP-PBL treatment lead to a transcriptomic profile that was more similar to that of cells in the steady-state (**Fig. 1A, S1**). These effects were not due to an impact of the postbiotic on cell viability (**Fig. S2**).

**Fig. 1:**
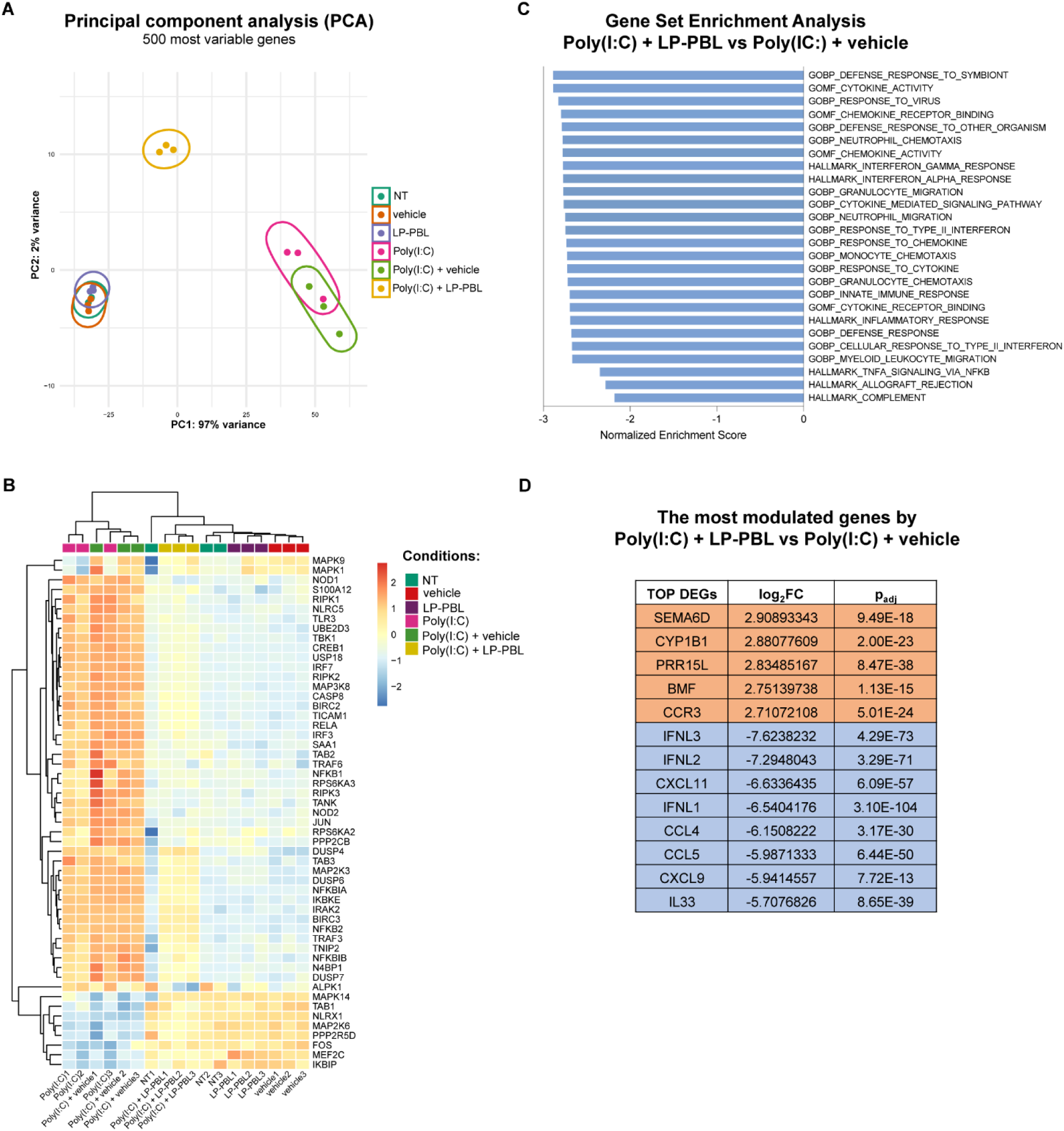
LP-PBL treatment shows anti-inflammatory signature on Poly(I:C)-stimulated keratinocytes. HaCaT keratinocytes were stimulated with 1 µg/ml Poly(I:C) and treated with 0.1 or 1 mg/ml LP-PBL or vehicle for 6 hours prior to RNA extraction and sequencing. (A) Principal component analysis (PCA) of the 500 most variable genes. (B) Heatmap representing the modulation of genes involved in TLR3 cascade by untreated cells (NT), vehicle, LP-PBL, Poly(I:C), Poly(I:C) + vehicle and Poly(I:C) + LP-PBL groups. (C) Graph representing the normalized enrichment score (NES) of negatively correlated gene sets identified in Poly(I:C) + LP-PBL group when compared to Poly(I:C) + vehicle and ranked by log2Fold Change. (D) Table representing the top 5 upregulated genes (in orange) (threshold > 2.7 log_2_FC) and the top 10 downregulated genes (in blue) (threshold < −5.7 log_2_FC) upon treatment with Poly(I:C) + LP-PBL compared to Poly(I:C) + vehicle.

Accordingly, the principal component analysis (PCA) showed clusterization of the transcriptomic profiles of unstimulated cells, postbiotic or vehicle treatment, indicating that LP-PBL did not on its own induce major changes in cellular gene expression (**Fig. 1A**). Differently, clustering of RNA-Seq data revealed distinct separation of Poly(I:C)-stimulated cells treated or not with vehicle compared to those treated also with LP-PBL. The latter group clustered separately from any other treatment indicating a control of the response initiated by Poly(I:C). Indeed, they were more similar to unstimulated cells as shown by the unsupervised hierarchical clustering analysis performed on the expression of TLR3 cascade genes (**Fig. 1B**). To deepen our analysis, using Gene Set Enrichment Analysis (GSEA) software, we compared Poly(I:C)-stimulated cells treated with LP-PBL or the vehicle. We observed the downregulation of some pathways that were found to be correlated with the activation of inflammation and recruitment of different immune cells, suggesting a potential role of the postbiotic in counteracting inflammation (**Fig. 1C**). Several of the most modulated genes by LP-PBL treatment are expressed in inflamed skin (**Fig. 1D**). For instance, we found downregulated interferons lambda (IFN-λs), which have been associated to AD [26] and psoriasis, especially *IFN-λ1* [27]; and are elevated in correlation with disease severity [28]. Also, some chemokine genes involved in the pathogenesis of skin diseases are downregulated by LP-PBL such as *CCL5*, *CXCL9*, *CXCL11,* as well as *CCL4* which plays a central role in the recruitment of pro-inflammatory cells [29–31]. In particular, *CCL4* level, which are high not only in AD patients’ blood [32] but also in acute AD lesions [29] and in psoriatic patients’ skin [29], is downregulated by LP-PBL. *IL-33* expression, instead, is upregulated mostly in AD [33] and is affected by LP-PBL concomitant treatment.

By exploring the most upregulated genes by LP-PBL treatment, we found that reduction of expression in some of them has been described in AD. For instance, *CYP1B1* is decreased in AD lesions and is involved in the aryl hydrocarbon receptor signaling pathway, which is dysregulated in AD patients [34]. *BMF* gene, which encodes for Bcl2 Modifying Factor, is involved in the regulation of apoptosis in AD [35] and its expression is reduced in AD patients’ eosinophils [36]. A similar effect could be occurring in keratinocytes and be counteracted by LP-PBL. Another gene upregulated by LP-PBL is *PRR15L*, gene encoding for proline-rich 15-like, whose function within keratinocytes is unknown, but AD skin shows reduced keratinocyte proline-rich proteins, which are involved in skin barrier dysfunctions and filaggrin regulation [37, 38].

Taken together, these results strongly suggest a protective role of LP-PBL on keratinocytes under inflammatory conditions by switching off the inflammatory response and the potential recruitment of immune cells, thus proposing an application of the postbiotic in skin disorders.

### LP-PBL treatment controls cytokine upregulation in keratinocytes under inflammatory conditions

Among the different cytokines known to be upregulated by Poly(I:C) binding to TLR3 [39–42] we found by RNA-seq that LP-PBL treatment counteracted the increase of *IL-6* [the 26^th^ most downregulated gene by LP-PBL in response to Poly(I:C) (log_2_FC −4,80 p_adj_ 3,98E-34)], *CCL-2* [311^th^ (log_2_FC −2,49 p_adj_ 5,07E-64) position] and *IL-8* [316^th^ (log_2_FC −2,47 p_adj_ 4,67E-31) position]. To validate the transcriptomic analysis, cells were stimulated with Poly(I:C) and treated with two concentrations of LP-PBL or vehicle. As expected, Poly(I:C)-stimulation increased the expression of *IL-6* (**Fig. 2A**)*, IL-8* (**Fig. 2B**), and *CCL-2* (**Fig. 2C**) in comparison to unstimulated cells. By contrast, LP-PBL treatment controlled pro-inflammatory cytokine expression induced by Poly(I:C) in a dose-dependent manner as compared to vehicle. In particular, we observed that LP-PBL was more effective in controlling *IL-6* upregulation compared to *IL-8* and *CCL-2*, thus validating the transcriptomic analysis. We then evaluated the effect of LP-PBL treatment at the protein level by ELISA. We observed that LP-PBL treatment on Poly(I:C)-stimulated cells could control even more effectively the protein levels of IL-6 (**Fig. 2D**), IL-8 (**Fig. 2E**), and CCL-2 (**Fig. 2F**) as compared to vehicle-treated cells.

**Fig. 2:**
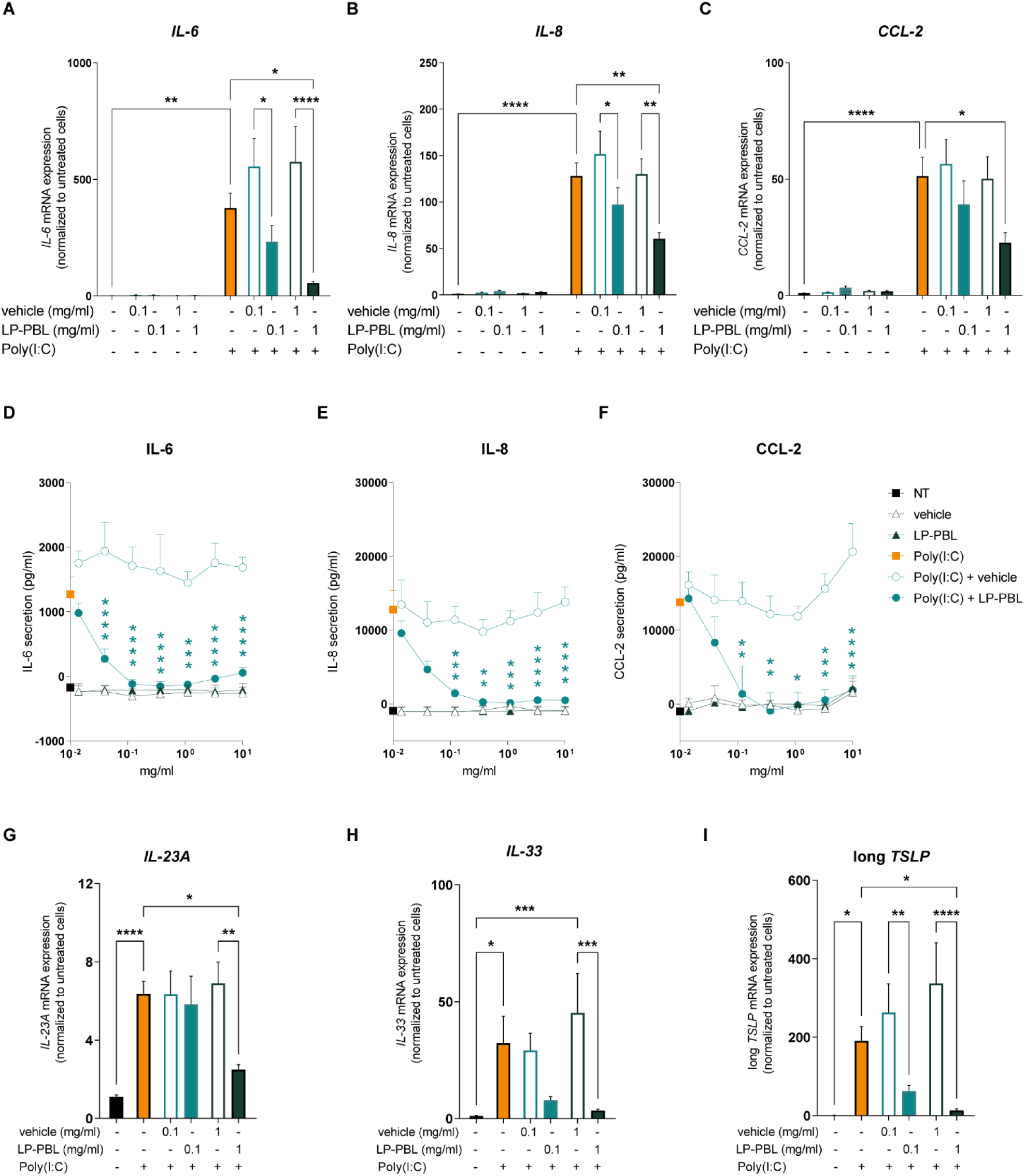
LP-PBL treatment controls cytokine upregulation in keratinocytes under inflammatory conditions. (A-C, G-I) HaCaT keratinocytes were treated for 6 hours with 0.1 or 1 mg/ml LP-PBL or vehicle following stimulation with 1 µg/ml Poly(I:C). mRNA expression levels of *IL-6* (A), *IL-8* (B), *CCL-2* (C), *IL-23A* (G), *IL-33* (H), and long *TSLP* (I) were analyzed by RT-PCR. (D-F) HaCaT keratinocytes were treated for 24 hours with different concentrations of LP-PBL or vehicle and/or stimulated with 1 µg/ml Poly(I:C). Release of IL-6 (D), IL-8 (E), and CCL-2 (F) was measured by ELISA. Each graph represents the mean ± SEM of three (A-C, G-I) and two (D-F) independent experiments. *Statistical analysis by one-way ANOVA, Tukey’s multiple comparisons test. Only comparisons between NT and Poly(I:C)-treated samples are represented (A-C, G-I), and two-way ANOVA between Poly(I:C) + vehicle and Poly(I:C) + LP-PBL, Šidák multiple comparisons test (D-F).

These results suggest that the postbiotic might be involved in the transcriptional and post-transcriptional regulation of these proteins. Considering the anti-inflammatory effect of LP-PBL and the strong reduction of inflammatory markers involved in psoriasis and AD, including *IL-33,* observed by transcriptomics (**Fig. 1D**), we next extended this analysis to evaluate whether the postbiotic could also counteract inflammatory skin mediators typical of psoriasis and AD inflamed skin.

Psoriatic skin is characterized by increased levels of IL-23 [43], particularly of its subunit IL-23A [44]; while AD patients’ skin is usually associated with the overexpression of cytokines like IL-33 and long TSLP [33, 45]. Transcriptomic analysis showed that LP-PBL was more efficient in controlling *IL-33* and long *TSLP* expression, the 10^th^ and 80^th^ most downregulated genes, respectively, than *IL-23A,* which is in the 1041^th^ position. Similarly, RT-PCR gene expression analysis showed that, after 6 hours of LP-PBL treatment, the postbiotic was able to reduce the Poly(I:C)-induced upregulation of *IL-23A* (**Fig. 2G**), and almost prevented the expression of *IL-33* (**Fig. 2H**), and long *TSLP* (**Fig. 2I**) in comparison to the vehicle in a dose dependent fashion.

Taken together, these results confirm the anti-inflammatory effect of LP-PBL treatment on Poly(I:C)-stimulated keratinocytes suggesting a potential therapeutic role in skin inflammation.

### LP-PBL treatment shows soothing effect on volunteers’ irritated skin

Considering the observed *in vitro* anti-inflammatory activity of LP-PBL, we decided to evaluate its effects in a clinical test on healthy volunteers using a model of skin irritation induced by sodium lauryl sulfate (SLS) (**Fig. 3A**). When SLS comes into contact with the skin, it disrupts the skin barrier, leading to increased water loss, dryness, and inflammation [46]. The subjects applied on their forearm patches containing 2% SLS aqueous solution, and after 24 hours, it significantly increased skin redness (**Fig. 3B**) and TEWL (**Fig. 3C**).

**Fig. 3:**
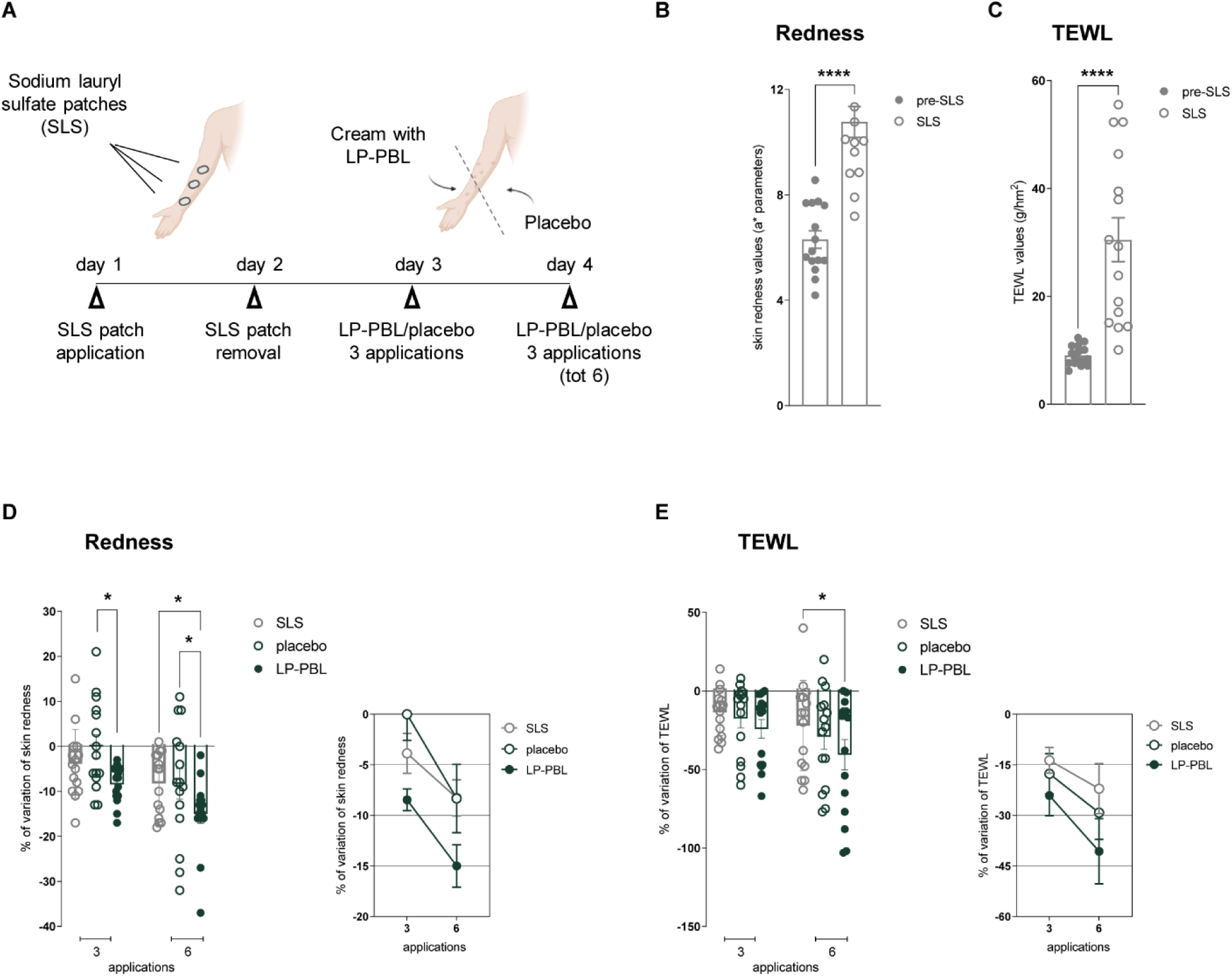
LP-PBL treatment shows soothing effect on volunteers’ irritated skin. (A) Schematic representation of experimental schedule: 15 healthy Caucasian volunteers’ forearm skin was irritated by occlusive patches containing 2% SLS aqueous solution for 24h. After 24 hours from patches removal, creams containing 2% LP-PBL or placebo were applied three times a day for 2 days. Instrumental evaluation of skin redness (B) and TEWL (C) before and after 24 hours of SLS patches application: quantification of skin redness (D) and TEWL (E) of LP-PBL efficacy in comparison to placebo after three and six applications. *Statistical analysis by paired t-test (B, C) and one-way ANOVA, Tukey’s multiple comparisons test (D, E).

After patches removal, volunteers applied on one side a cream containing LP-PBL and, on the other, a placebo cream (the same formulation but without LP-PBL) three times a day for two days. Following three and six consecutive applications, the postbiotic-containing cream significantly decreased skin redness compared to placebo (**Fig. 3D**) and reduced skin TEWL induced by SLS (**Fig. 3E**). These findings confirm the anti-inflammatory properties of LP-PBL treatment demonstrated *in vitro* and show a remarkable and rapid restorative effect on the irritated skin.

### LP-PBL treatment preserves barrier integrity in keratinocytes

The capacity of the postbiotic-containing formulation to counteract the increase in TEWL observed in the clinical study prompted us to evaluate the effect of LP-PBL in keratinocytes barrier properties. Increased TEWL levels have been linked to decreased expression of filaggrin, one of the principal epidermal differentiation-related molecules in keratinocytes. Reduced levels of filaggrin and consequently its metabolites, including NMFs, increase skin pH that induces damages on the barrier and increases the TEWL [47]. Furthermore, defects in TJ proteins contribute to barrier abnormalities that characterize AD skin [3, 48]. Hence, we investigated the effect of LP-PBL on keratinocytes barrier-related proteins.

We observed that LP-PBL dose-dependently upregulated *FLG* (**Fig. 4A**) and *ZO-1* expression (**Fig. 4D**) in Poly(I:C)-stimulated keratinocytes compared to vehicle-treated cells. Immunofluorescence staining (**Figs. 4B, E**) confirmed these data at the protein level and revealed that LP-PBL treatment significantly increased filaggrin levels (**Fig. 4C**) and preserved ZO-1 integrity (**Fig. 4F**) compared to Poly(I:C)-vehicle-treated cells.

**Fig. 4:**
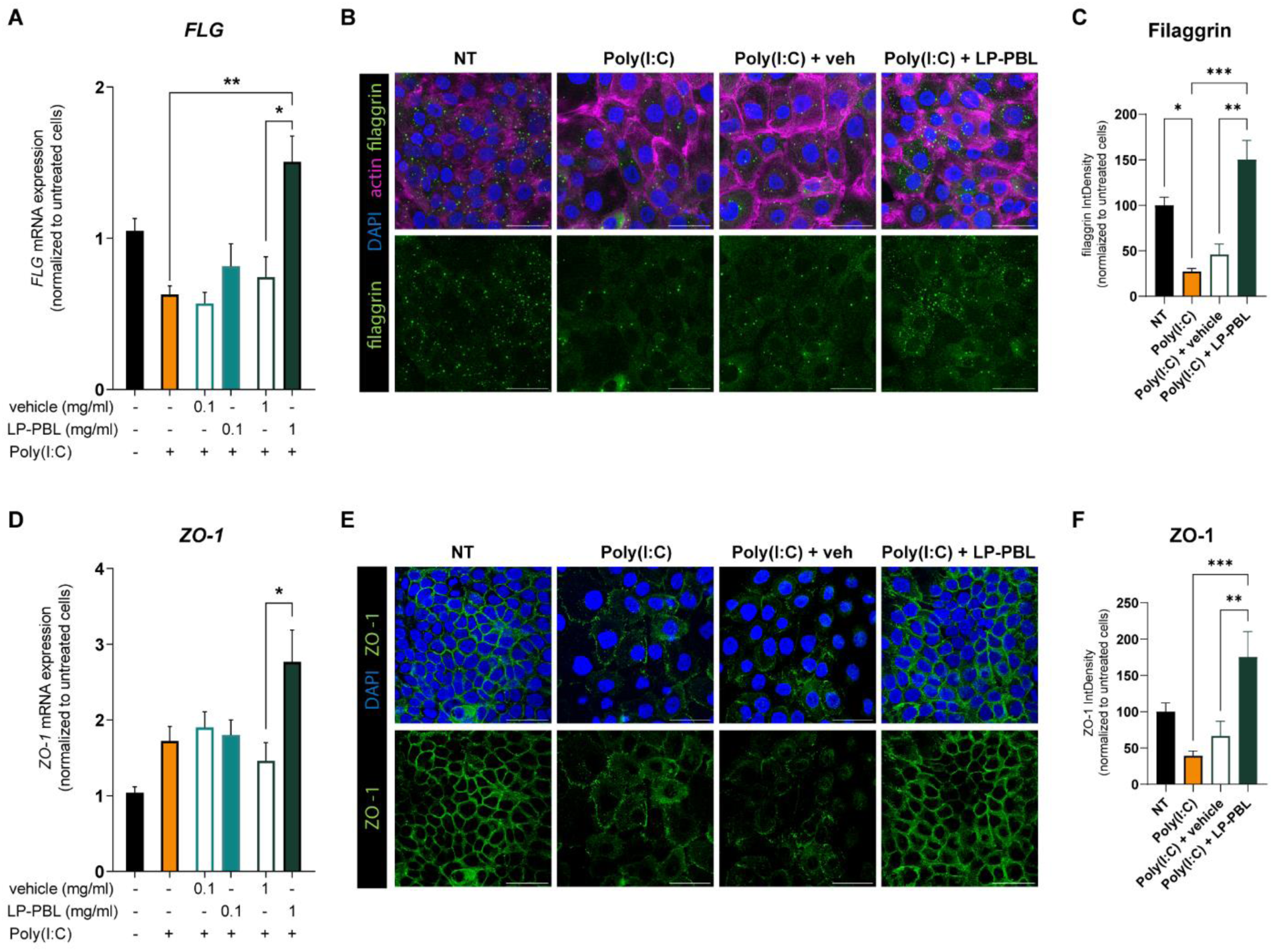
LP-PBL treatment preserves barrier integrity on Poly(I:C)-stimulated keratinocytes. HaCaT keratinocytes were treated for 6 hours with 0.1 or 1 mg/ml LP-PBL or vehicle following stimulation with 1 µg/ml Poly(I:C). mRNA expression levels of *FLG* (A) and *ZO-1* (D) were analyzed by RT-PCR. (B, E) HaCaT keratinocytes were stimulated with 1 µg/ml Poly(I:C) and treated for 24 hours with 1 mg/ml LP-PBL or vehicle. Analysis of filaggrin (C) and ZO-1 (F) was performed by immunofluorescence. Scale bar – 50 µm. Objective 63X. Each graph represents the mean ± SEM of three (A, D) and two (C, F) independent experiments. *Statistical analysis by one-way ANOVA, Tukey’s multiple comparisons test.

Taken together, these data strongly suggest that the postbiotic treatment can effectively preserve skin barrier integrity in keratinocytes under inflammatory conditions, highlighting the protective role of LP-PBL in skin barrier functions.

## DISCUSSION

The skin protects our body from mechanical damage and external factors by acting as a shield against invasion of pathogens and antigens. It also plays a key role in the maintenance of thermal regulation and control of body water loss. In particular, keratinocytes in the outer layer of skin prevent excessive TEWL and the increase in barrier permeability, but also protect against ultraviolet radiations and participate in the immune response [1]. For these reasons, the preservation of skin barrier integrity is essential to prevent tissue injuries and maintain a proper skin homeostasis.

Barrier damage as well as abnormal immune response within the skin are among the factors contributing to barrier dysfunctions and leading to inflammatory diseases like psoriasis and AD [49, 50]. The pathogenesis of these common chronic skin disorders is multifactorial and includes immune dysregulation, barrier damage, exposure to environmental allergens, skin dysbiosis, and pathobiont overgrowth [4, 5, 51, 52].

Nowadays, there is a growing interest in alternative and complementary strategies to treat chronic inflammatory skin diseases. Considering the central role of the skin microbiota in the onset of these diseases, topical application of skin commensal bacteria has been proposed [10], in particular to ameliorate AD signs such as inflammation, epithelial barrier impairments, skin redness, pruritus, and also to reduce *Staphylococcus aureus* abundance, typical of AD skin [53].

However, topical application of probiotics is not exempt from side effects like the possibility of antibiotic resistance [11] and potential translocation of probiotics across damaged skin. The use of probiotics is indeed not recommended in some pathological conditions, such as inflammatory bowel disease, as during the acute phase of inflammation, due to the lack of protective barriers, the use of probiotics may even exacerbate the condition [14]. Further, the presence of antimicrobial preservatives added in formulations can influence the viability of both probiotics and commensal bacteria [54], while additives are required to ensure probiotic stabilization because of their sensitivity to temperature, humidity levels, pH, and osmolarity [10].

Postbiotics can represent a safe and innovative alternative to probiotics. Their definition remains a controversial issue in the scientific community but, because of the instability of dead bacteria or their fragments, we define postbiotics as “any factor resulting from the metabolic activity of a probiotic or any released molecule capable of conferring beneficial effects to the host in a direct or indirect way” [18–20]. For this reason, they are not as sensitive to temperature and humidity levels as bacteria, and therefore show several technical advantages over probiotics [22]. Moreover, it is known that postbiotics exhibit a variety of beneficial effects, including anti-inflammatory activities, modulation of immune response, and maintenance of intestinal epithelial barrier integrity [12, 18, 21, 22].

It is well established that keratinocytes in the epidermis play a central role in the activation of an immune response against external microbes. They express a variety of TLRs [55] activating different signaling pathways upon exposure to exogenous ligands. In particular, they express TLR3 that recognizes dsRNA, associated with viral infection, and activates a series of intracellular signaling events within keratinocytes including pro-inflammatory molecules release [2]. Moreover, TLR3 signaling is involved in barrier dysfunction and it has been found overexpressed in AD and psoriatic skin [56, 57]. Several studies have shown that Poly(I:C)-stimulation induces in keratinocytes high levels of IL-6 [41], IL-23 [58] and IL-8 [39] as well as chemokine (C-C motif) ligand 2 (CCL-2) [40], that recruits immune cells, intensifying the inflammatory response. It has also been shown that these inflammatory mediators are overexpressed in psoriatic skin [59–62].

Therefore, we questioned if the postbiotic treatment could modulate the inflammatory response upon stimulation with the TLR3 agonist, Poly(I:C). We observed that LP-PBL treatment inhibited the upregulation and release of inflammatory mediators, such as IL-6, IL-8, and CCL-2 suggesting its role in counteracting the inflammatory process.

Given the promising anti-inflammatory effect of LP-PBL, we decided to evaluate this activity in a clinical test on healthy volunteers’ skin previously irritated by SLS, which leads to red, dry, and irritated skin [46]. We observed that topical treatment of LP-PBL versus placebo reduced skin redness and TEWL highlighting LP-PBL capacity to induce a fast recovery of subjects’ irritated skin.

We observed that LP-PBL was able to preserve keratinocyte barrier properties which are normally sustained by the presence of proteins like filaggrin and TJs. Filaggrin is a crucial protein in keratinocytes with pleiotropic functions, and mutations in the *FLG* gene represent the strongest genetic risk factor for AD barrier dysfunction [47]. Reduction in filaggrin expression has been associated to impairment of epidermal proteins organization, abnormalities in lipid production and increase in skin pH, features linked to skin dryness and increased TEWL [3, 47].

Filaggrin deficiency in AD skin is often associated with impairment in TJ proteins, including ZO-1 [50, 63]. TJs are key structures that actively contribute to the maintenance of skin barrier integrity. Indeed, they connect keratinocytes and regulate the passage of allergens and microbes as well as water and ions, thus contributing to prevent excessive TEWL [50]. We showed that LP-PBL treatment successfully increased ZO-1 and filaggrin levels in Poly(I:C)-stimulated cells, highlighting its potential protective role on skin barriers.

We performed RNA-seq on keratinocytes under inflammatory conditions and we observed that LP-PBL treatment modulated some pathways related to the activation of inflammation and recruitment of different immune cells. In particular, among the most downregulated genes we found many that are involved in skin inflammatory disorders, like *CCL4, CCL5, CXCL9,* and *CXCL11* that are overexpressed in psoriatic lesions, while modulation of *CYP1B1*, *BMF*, *IFN-λ1*, and *IL-33* are implicated in AD. These findings suggest that LP-PBL could not only preserve barrier integrity, but also counteract the inflammation typical of these skin disorders.

We showed that LP-PBL treatment controlled the upregulation of the long isoform of *TSLP,* which is known to stimulate dendritic cells to produce Th-2-attracting chemokines and initiate Th-2 responses. Indeed, we have shown that long TSLP is undetectable in healthy skin and upregulated in many Th-2-related diseases including AD [64]. We also showed that LP-PBL was able to modulate *IL-23A* and *IL-33*, usually overexpressed in psoriatic and AD skin, respectively [44, 65]. IL-23A and IL-33 neutralizing antibodies have been approved as treatment for these disorders [66, 67], but they do not counteract the cause of the inflammation. By contrast, the evidence that LP-PBL re-established the barrier properties of the keratinocytes and controlled the upregulation of these cytokines is an encouraging outcome, suggesting that LP-PBL might be a complementary therapeutic treatment for these patients, and may prolong that time between flares or even abrogate them. Furthermore, our results are complemented by an international clinical study that has recently demonstrated the beneficial effects of a cream formulation containing LP-PBL and prebiotics on AD skin [68]. The study was based on 396 subjects of both sexes divided in three age groups (infants, children, adults) suffering from mild-to-moderate AD. The patients applied on their eczema regions a cream formulation containing LP-PBL for 15 weeks. In line with a decrease in SCORAD, PRURISCORE, and IGA scores, the treatment showed a good efficacy in reducing the severity of AD, already after the first month of treatment. LP-PBL-based-cream also reduced the number of episodes that require topical treatment with corticosteroids. Importantly, during the treatment, patients reported an improvement in their quality of life, which is known to be compromised in AD [68].

## CONCLUSION

In conclusion, here we showed that treatment with LP-PBL, a *Lactobacillus paracasei* CNCM I-5220-derived postbiotic, effectively mitigates skin inflammation and preserves skin barrier integrity. In particular, LP-PBL could control in keratinocytes the upregulation of several interleukins and chemokines involved in AD as well as psoriasis pathogenesis.

Supported by the findings of Gelmetti and colleagues [68], the results of our study suggest that topical application of LP-PBL could offer a safe and promising new approach for managing skin inflammation and counteracting barrier dysfunctions across various skin inflammatory conditions, including psoriasis.

## AUTHOR CONTRIBUTION

SP and FA designed, performed, and analyzed the experiments. NT and DB performed all the bioinformatic analyses. SP, FA, GP and MR wrote the manuscript. SP, FA, NT, DB, GP and MR revised the manuscript. All authors contributed to the article and approved the submitted version.

## FUNDINGS

This work was supported by Postbiotica S.r.l.

## CONFLICT OF INTEREST

MR is co-founder and CSO of Postbiotica S.r.l. GP is co-founder and scientific advisor of Postbiotica S.r.l. FA and NT are employees of Postbiotica S.r.l. The remaining authors declare that the research was conducted in the absence of any commercial or financial relationships that could be construed as a potential conflict of interest.

---

**Fig. S1:**
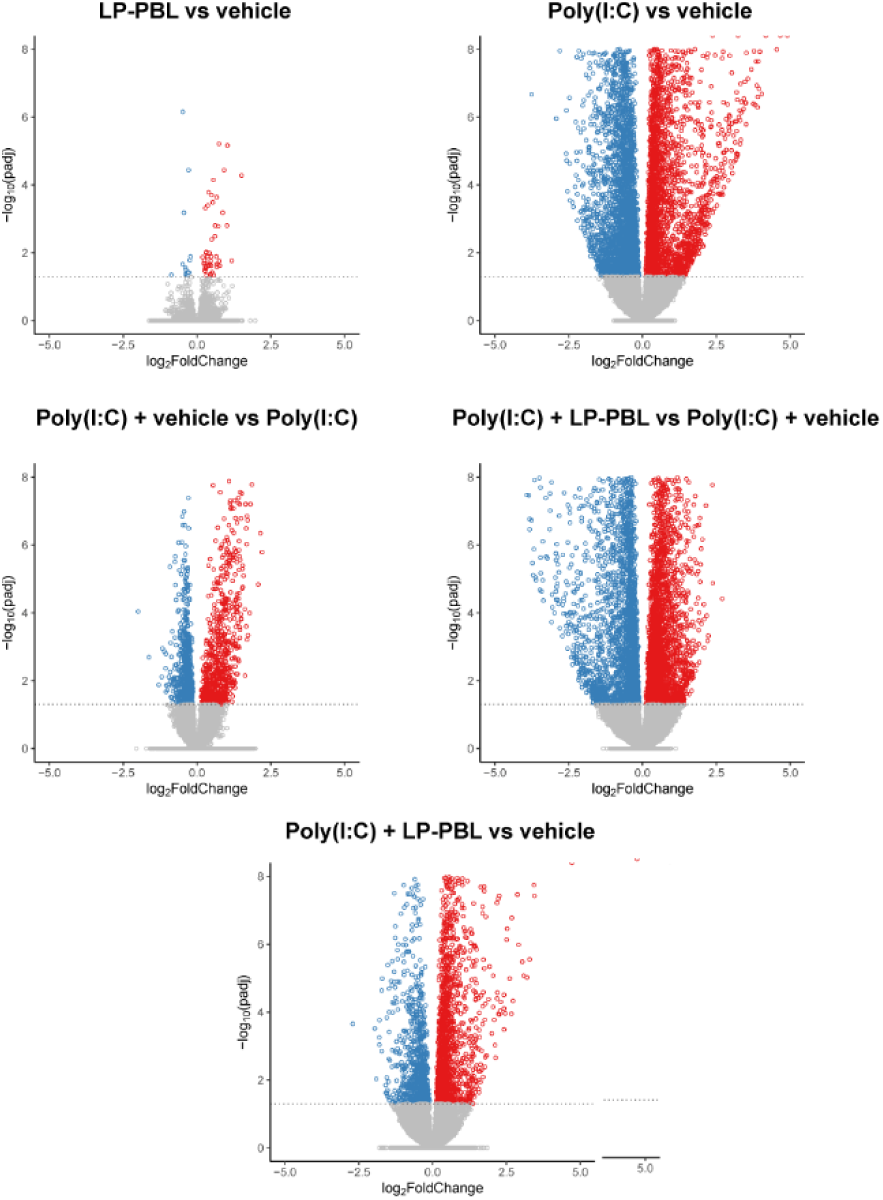
Volcano plots representing the number of genes upregulated (in red) or downregulated (in blue): LP-PBL treatment in comparison to the vehicle in steady-state (top left panel); Poly(I:C)-treated cells in comparison to unstimulated cells (NT) (top right panel). Comparison between genes modulated by Poly(I:C) + vehicle vs Poly(I:C) group (middle left panel), Poly(I:C) + LP-PBL vs Poly(I:C) + vehicle (middle right panel) and Poly(I:C) + LP-PBL vs vehicle (bottom panel) (padj < 0.05).

**Fig. S2:**
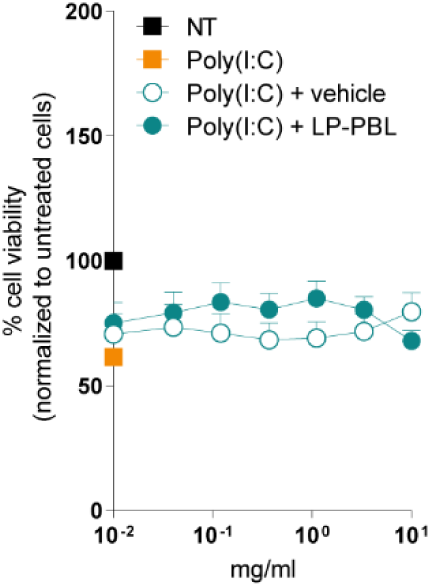
HaCaT keratinocytes were treated with different concentrations of LP-PBL or vehicle following 1 µg/ml Poly(I:C)-stimulation. After 24 hours MTT assay was performed. Results are presented as the percentage of cellular viability of untreated cells. The graph represents the mean ± SEM of two independent experiments. *Statistical analysis by two-way ANOVA, Šidák multiple comparisons test, between Poly(I:C) + LP-PBL and Poly(I:C) + vehicle.

